# Early innate immune signatures correlate with Ad26.COV2.S vaccine durability and protective efficacy

**DOI:** 10.64898/2026.06.09.731069

**Authors:** Alessandro Colarusso, Kathryn E. Stephenson, Ai-Ris Y. Collier, Frank Wegman, Roland Zahn, Malika Aid, Dan H. Barouch

## Abstract

The innate immune system is rapidly activated following antigen exposure and plays a critical role in shaping the ensuing adaptive immune responses. In this study, we performed bulk RNA sequencing and proteomic profiling in a cohort of 25 rhesus macaques following Ad26.COV2.S immunization to characterize early innate correlates of vaccine immunogenicity and protection. Our results show that innate immune activation occurred as early as day 1 post-vaccination and demonstrated an systemic enrichment of antiviral interferon pathways, interleukin signaling, and innate immune cell signatures. These early transcriptomic signatures correlated positively with humoral and cellular immune responses at 6 weeks following vaccination and correlated inversely with viral loads following SARS-CoV-2 challenge. Similar correlates of immunogenicity were observed in a cohort of 25 adult healthy participants vaccinated with Ad26.COV2.S. Taken together, these findings highlight the importance of early activation of the innate immune system for Ad26.COV2.S vaccine immunogenicity and protective efficacy.

**Graphical Summary:** Graphical summary of the model proposed by our findings. Day 1 innate immune signatures are protective of long-term protection.

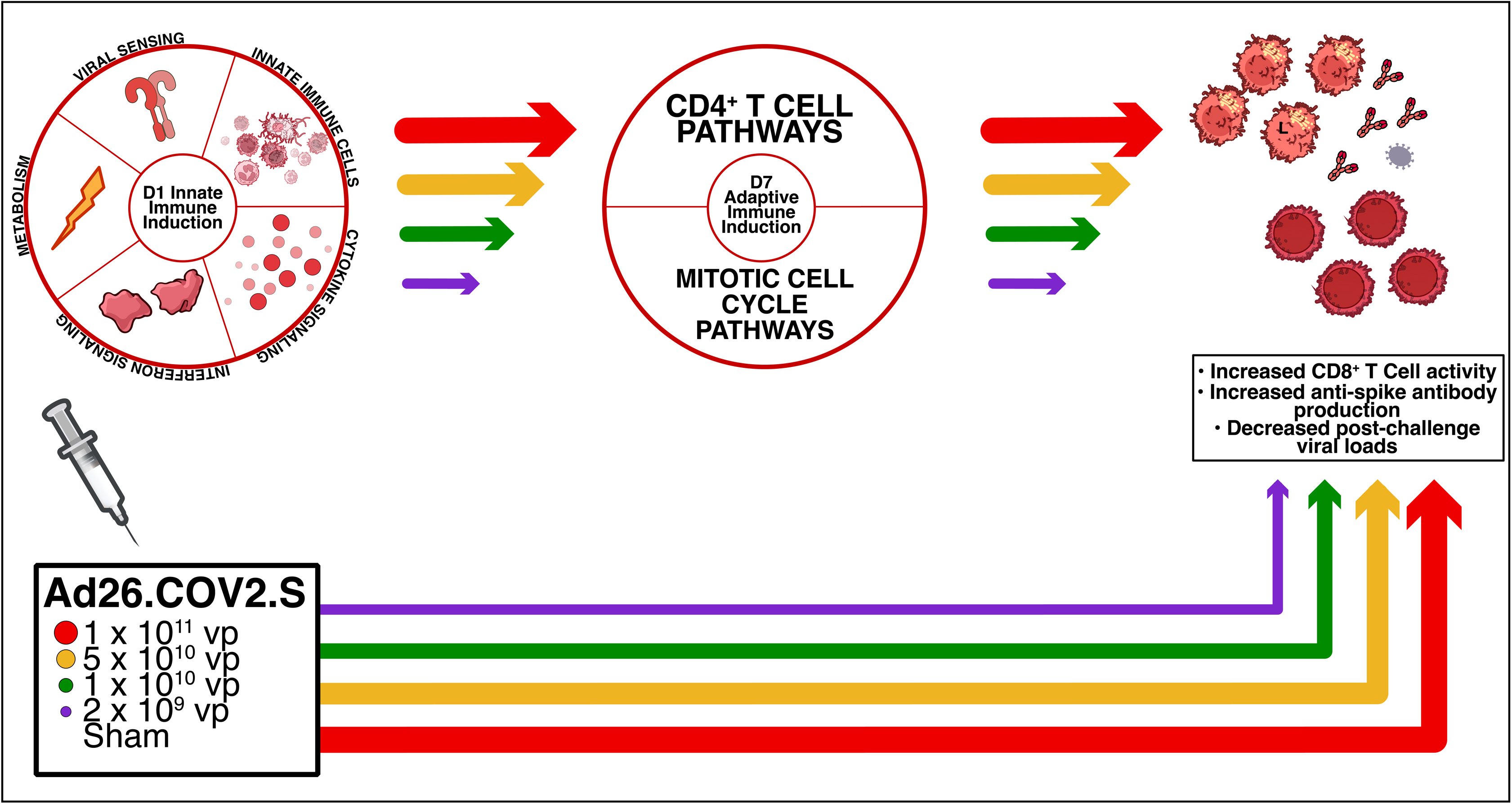

**IMPORTANCE:** Early vaccine induction of innate immunity may be critical for vaccine immunogenicity and protective efficacy. In this study, we evaluated innate immune responses following Ad26.COV2.S vaccination in both rhesus macaques and humans. Early induction of innate immune signatures on day 1 following vaccination correlated with subsequent development of adaptive immune responses. These data suggest that the immune programming that occurs immediately after vaccination dictates immunogenicity weeks or months later.

## INTRODUCTION

Innate immunity serves as the initial rapid host response to pathogens and typically precedes the induction of adaptive immune response by weeks or months^1,2^. Understanding the interaction between innate and adaptive immunity is crucial for effective vaccine design^3^. Adenoviral vectors have been widely utilized as vaccine platforms, including an adenoviral serotype 26 (Ad26) vaccine expressing SARS-CoV-2 (Ad26.COV2.S), which was developed by Johnson & Johnson and was authorized for emergency use^4^. The clinical efficacy^5^ and immunogenicity^6^ of Ad26.COV2.S have been reported in detail^7, 8^, yet the mechanisms by which Ad26 vaccination shapes early innate immune responses remain poorly understood^9^.

In this study, we evaluated early multi-omic signatures induced by Ad26.COV2.S and to determine if these signatures correlated with adaptive immune responses and protective efficacy. We used transcriptomic and proteomic profiling to define early innate immune pathways in 25 rhesus macaques vaccinated with Ad26.COV2.S in a previously published study^10^ (**Figure 1A**). We also performed similar multi-omic profiling in 25 humans who participated in an Ad26.COV2.S clinical trial (NCT04436276)^7^ (**Figure 1B**). Participants received an initial immunization of Ad26.COV2.S of either 1 x 10^11^ viral particles (vp), 5 x 10^11^ vp or placebo. Biological samples for multi-omic assays were collected on day 1 and day 7 following initial immunization. Additionally, some participants received a follow-up immunization (boost) at day 56 following initial immunization, and additional serological samples were collected^7^.

**Figure 1:**
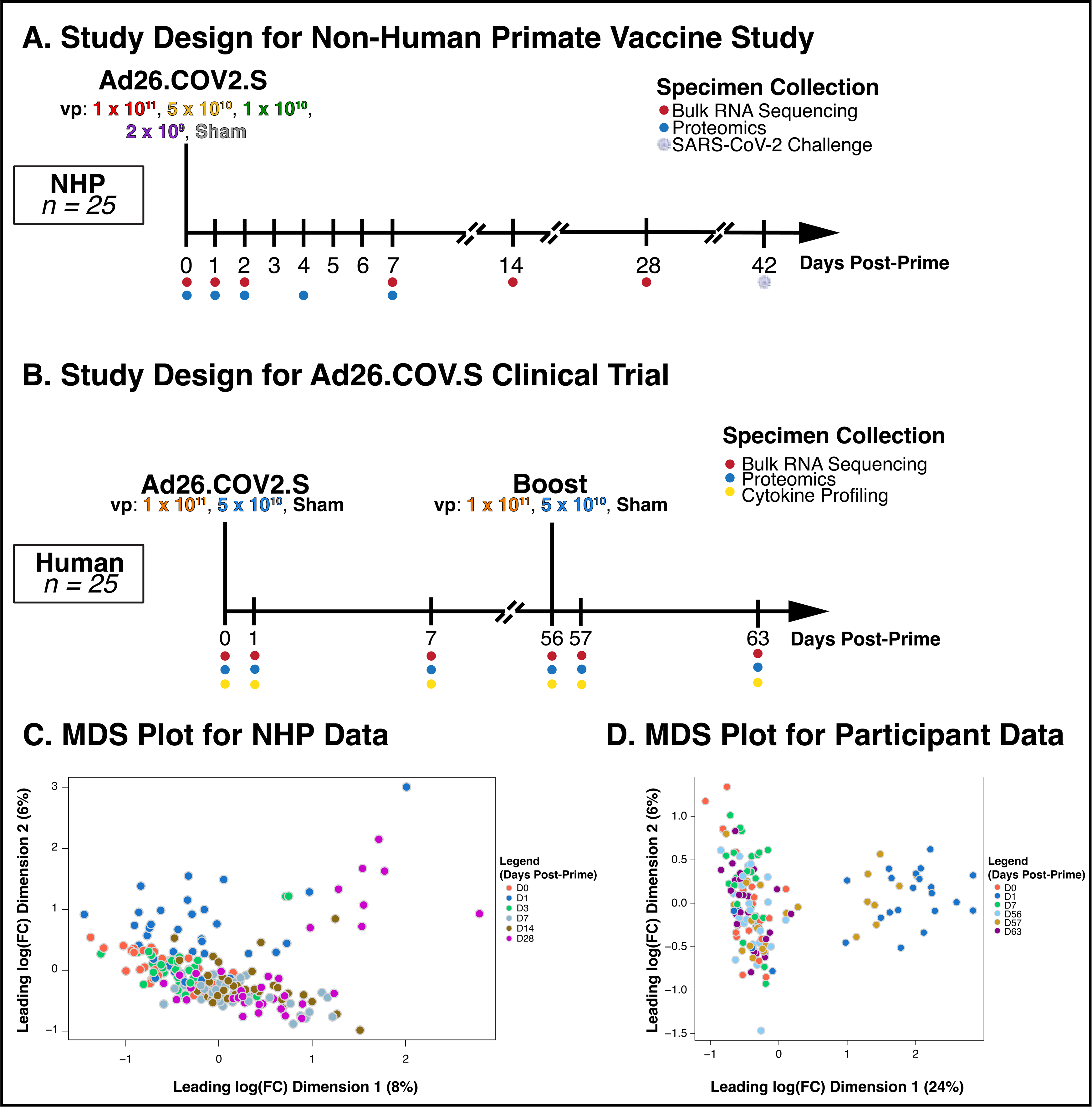
Immune kinetics induced by an Ad26.COV2.S using multi-omic profiling. **A.** Macaques (*N* = 5 per dose group) were immunized at baseline, and longitudinal blood samples were collected for bulk RNA sequencing and proteomics analysis. Animals were challenged with SARS-CoV-2 at day-42 following initial immunization. **B.** Participants (*N* = 10 per dose group) in a clinical trial (NCT04436276) were immunized at baseline, and longitudinal blood samples were collected for bulk RNA sequencing and proteomics analysis. Participants received a booster dose (*N* = 5 per group) at day-56 following initial immunization. **C.** Multidimensional scaling plot (MDS) for bulk RNA sequencing data collected in whole blood from rhesus macaques (*N* = 5 per group). **D.** MDS plot for bulk RNA sequencing data collected in whole blood from participants (*N* = 10 per group).

In this study, we quantified early innate immune signatures induced by Ad26.COV2.S vaccination in both macaques and human participants, with an emphasis on interferon mediated as well as innate immune cellular pathways. We evaluated these pathways and their relationships to subsequent adaptive immune responses and protective outcomes.

## RESULTS

### Innate immune pathways were upregulated by day 1 and T cell pathways were upregulated by day 7 following Ad26.COV2.S immunization of rhesus macaques

We investigated the early innate immune profiles following Ad26.COV2.S immunization in rhesus macaques. We immunized five groups of macaques (N = 5/group) with Ad26.COV2.S at doses of 1 x 10^11^, 5 x 10^10^, 1 x 10^10^, and 2 x 10^9^ vp, and a sham group^10^. The macaques were then challenged with SARS-CoV-2 at week 6 following vaccination. Blood samples were collected at baseline prior to immunization and on days 1, 3, 7, 14, and 28 following vaccination. We performed whole-blood transcriptomics to assess longitudinal transcriptomic signatures (**Figure 1A**)^8, 10^. A multidimensional scaling (MDS) plot revealed clustering for day 1 samples compared to other timepoints (**Figure 1C**)^11^. Using the high dose group for our discovery cohort, we performed Gene Set Enrichment Analysis (GSEA)^12, 13^ to gauge pathway activity following immunization as compared to baseline. Only pathways that were deemed significant by GSEA, defined as having a False Discovery Rate (FDR) ≤ 0.05^12^, were considered for downstream analysis.

To ensure we evaluated only vaccine-induced pathways, we performed GSEA on the sham group between day 1 following immunization and baseline: we excluded pathways that were significantly upregulated in the sham group on day 1. Similarly, we performed GSEA at baseline between vaccinated groups and the sham group. We did not consider pathways that were significantly upregulated in the vaccinated groups at baseline for further analysis. A complete list of the GSEA analysis of pathways is shown in **Supplementary Table 1**.

Major pathway modules identified on day 1 following immunization included antiviral interferon pathways, dendritic cell and monocyte pathways, and viral sensing pathways such as Toll-like receptor (TLR) pathways (**Figure 2**). In addition, inflammatory pathways such as chemokine and complement activity were significantly upregulated on day 1. Consistent with the rapid kinetics in mice^14^, many innate immune pathways that were activated by day 1 were largely resolved by day 3 (**Figure 2**). We observed — via Normalized Enrichment Score (NES)^12^ — a dose-dependent upregulation of interferon and proinflammatory signaling pathways, consistent with our previous work^8^. Interferon activity pathways (*ANTIVIRAL IFN SIGNATURE*, NES = 2.47, FDR = 2 x 10^−4^) and monocyte activity (*ENRICHED IN MONOCYTES*, NES = 2.41, FDR = 2 x 10^−4^) were significantly higher in the high-dose group (**Figure 2**).

**Figure 2:**
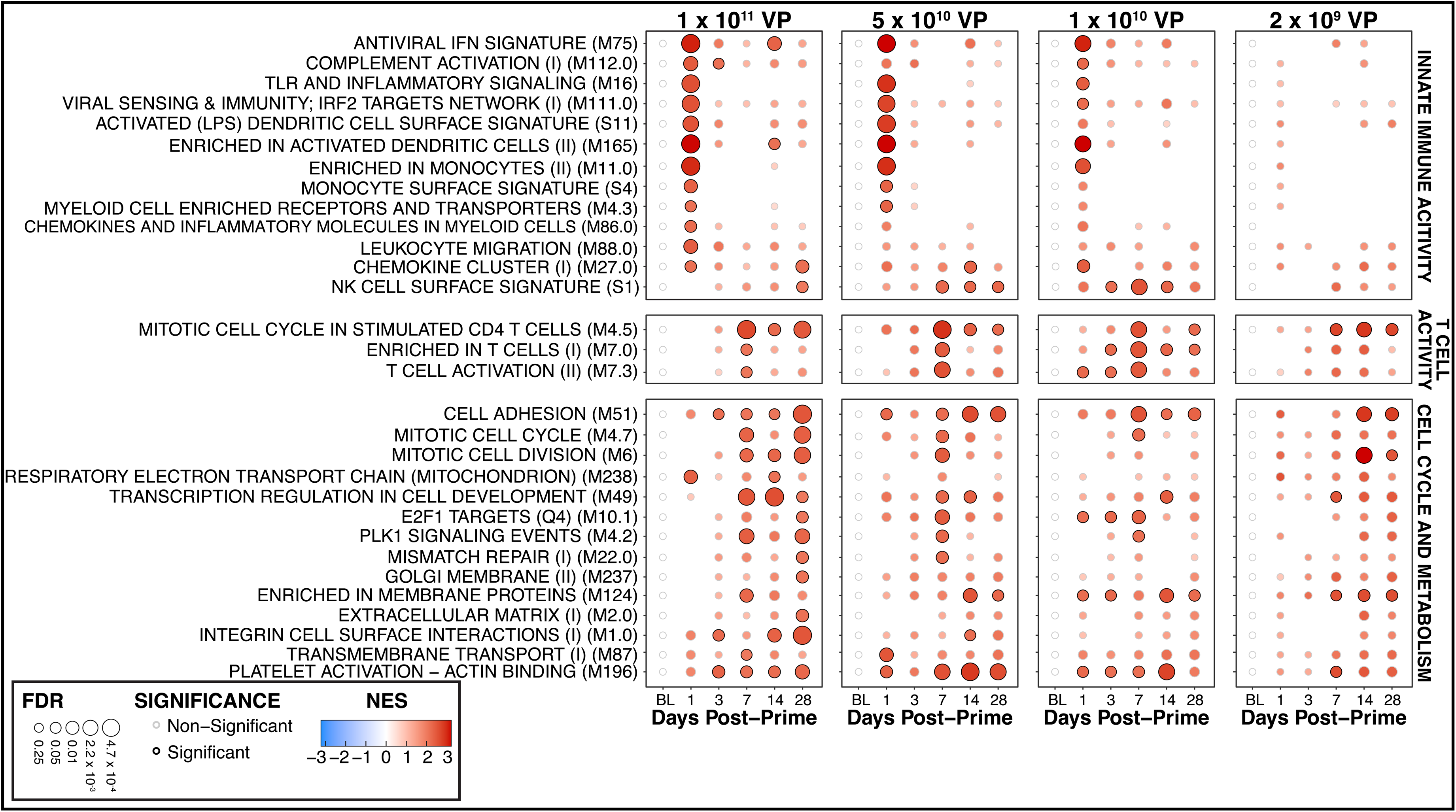
Ad26 vector immunization stimulates rapid activation and resolution innate immune pathways in rhesus macaques within one day following initial immunization. Similarly, T cell and associated cell proliferation pathways are activated at seven days following initial immunization. Transcriptomic pathway analysis reveals that the highest-dose group (1 x 10^11^ vp, *N* = 5) shows the most robust activation of pathways, with lower pathways exhibiting similar patterns of pathway activation at reduced magnitude and significance.

On day 7 following immunization, we identified significant upregulation of different pathways associated with T cell signaling (*ENRICHED IN T CELLS,* NES = 1.68, FDR = 0.03) and activity (*T CELL ACTIVATION (II),* NES = 1.66, FDR = 0.04) as well as cell metabolism (e.g. *MITOTIC CELL CYCLE,* NES = 1.87, FDR = 4 x 10^−3^) (**Figure 2**). These pathways were generally not increased on day 1. We observed an increase in T cell activation, cell adhesion, and mitotic cell cycle pathways, including classical mitosis regulatory genes such as *PLK1*^15^ and *E2F1*^16^. Specifically, we observed that the CD4^+^ T cell proliferation pathway (*MITOTIC CELL CYCLE IN STIMULATED CD4 T CELLS*, NES = 2.13, FDR = 2 x 10^−4^) was the most enriched and significant pathway in the T Cell Activity module (**Figure 2**). This pathway’s activity was reduced but still observed on day 14 (NES = 1.69, FDR = 0.04) and day 28^17^ (NES = 1.87, FDR = 2 x 10^−3^) (**Figure 2**).

Taken together, these data suggest upregulation of antiviral and acute inflammatory pathways on day 1 following immunization and upregulation of T cell, cell cycle, and metabolic pathways on day 7 following immunization.

### Innate pathways identified on day 1 following immunization correlated with transcriptomic pathways identified on day 7 following immunization

We performed pathway enrichment analysis for individual animals using single-sample Gene Set Enrichment Analysis (ssGSEA)^18^. We then performed Spearman rank correlations between these ssGSEA transcriptomic signatures in macaques on day 1 and day 7. We found that day 1 interferon, complement, dendritic cell pathways were the major modules that correlate positively with day 7 CD4+ T cell cycle and associated mitotic cell cycle pathways, with Spearman ρ coefficient ranging from 0.13 to 0.73, and most (70%) Bonferroni adjusted p-values, *p*_adj_ < 0.05. (**Figure 3, Supplementary Table 2**). Namely, the strongest and most significant correlation was observed to be between day 1 dendritic cell activity (*ENRICHED IN ACTIVATED DENDRITIC CELLS*) and day 7 PLK1 signaling (*PLK1 SIGNALING EVENTS*), with a Spearman coefficient ρ = 0.73, and Bonferroni adjusted p-value, *p*_adj_ = 3 x 10^−4^. This association of day 1 transcriptomic signatures to mitosis and cell proliferation at day 7 post-immunization suggested that day 1 innate immune signatures were predictive of day 7 T cell activity^19^.

**Figure 3:**
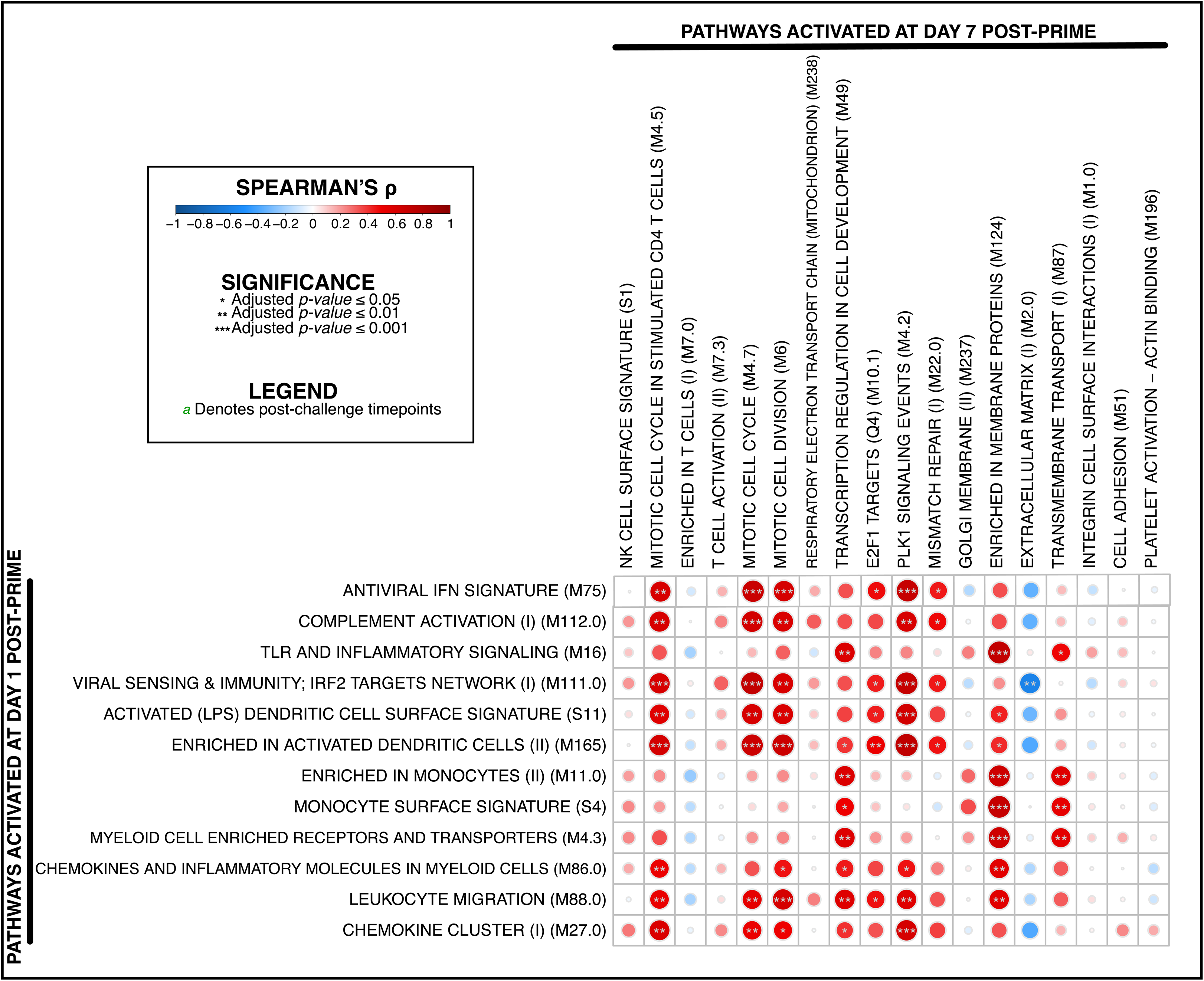
Whole-blood transcriptomic innate immune signatures at day 1 following initial Ad26.COV2.S immunization in macaques are predictive of T cell and cell proliferation signatures at day 7 following initial immunization, with an increase in vaccine cargo dose being predictive of more robust protection. *N* = 25 macaques per group. A Spearman-rank test was used to calculate correlations, and *p-*values were adjusted for multiple comparisons using a Bonferroni correction.

### Transcriptomic pathways on day 1 shaped adaptive immune responses

Antigen-specific immune responses are the hallmarks of vaccines^20^. We therefore sought to evaluate whether early transcriptomic signatures on day 1 following Ad26.COV2.S vaccination correlated with adaptive immune responses and protective efficacy in macaques. Using a Spearman-rank test, we correlated ssGSEA NES to previously reported spike RBD-specific binding antibody responses by enzyme-linked immunosorbent assays (ELISA) and pseudovirus neutralization assays on day 42^10^ (**Supplementary Table 3**). We observed that antiviral interferon signatures (0.53 ≤ ρ ≤ 0.80; 8 x 10^−6^ ≤ *p*_adj_ ≤ 3 x 10^−3^), viral sensing (0.54 ≤ ρ ≤ 0.78; 3.4 x 10^−6^ ≤ *p*_adj_ ≤ 5.5 x 10^−3^), dendritic cell signatures (0.38 ≤ ρ ≤ 0.76; 8.6 x 10^−6^ ≤ *p*_adj_ ≤ 5.8 x 10^−2^), and chemokine pathways^13^ (0.17 ≤ ρ ≤ 0.68; 1.6 x 10^−4^ ≤ *p*_adj_ ≤ 4.1 x 10^−1^) correlated positively with both ELISA and NT50 titers (**Figure 4**). These pathways also correlated with cellular immunity as measured by interferon-γ (IFN-γ) Enzyme-Linked ImmunoSpot (ELISpot) assays (0.14 ≤ ρ ≤ 0.67; 2.3 x 10^−4^ ≤ *p*_adj_ ≤ 5.0 x 10^−1^) (**Figure 4**).

**Figure 4:**
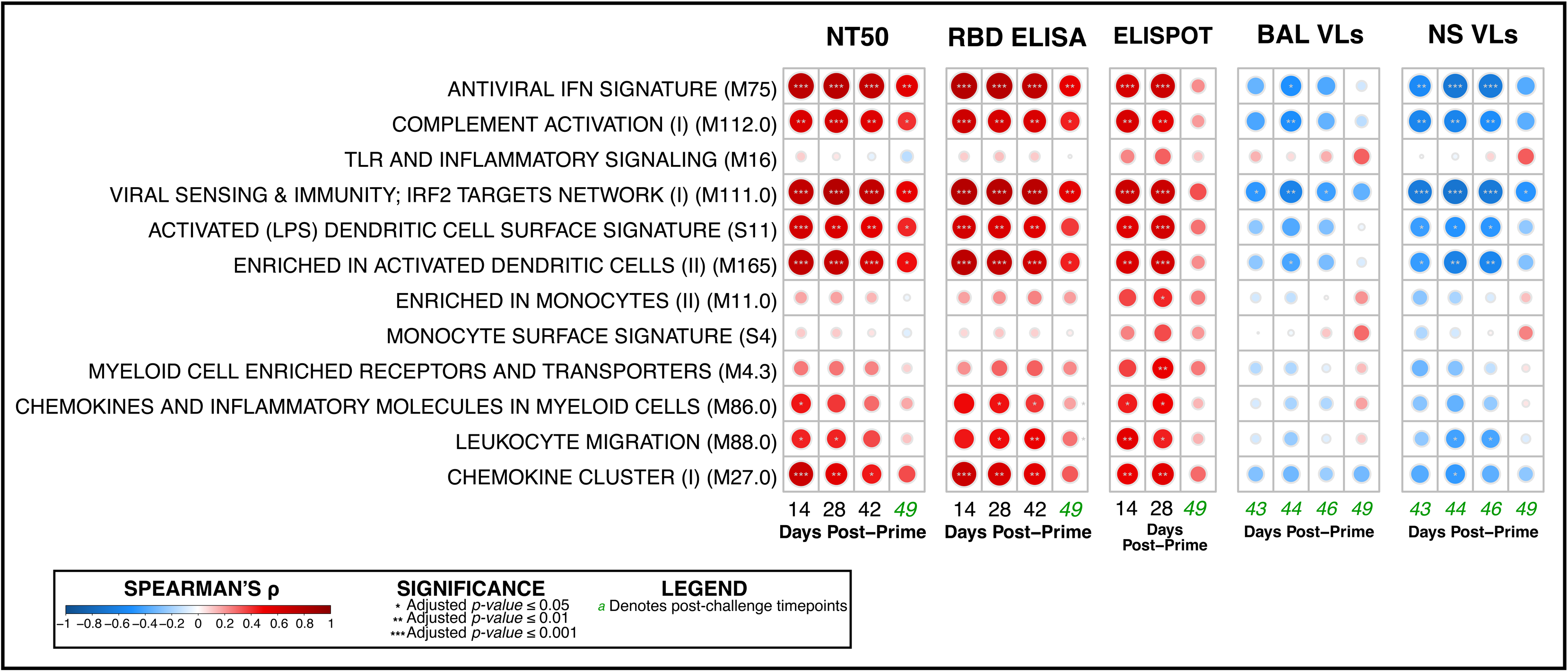
Whole-blood innate immune transcriptomic signatures at day 1 following initial Ad26.COV2.S immunization in macaques are predictive of antibody and T cell responses at two to seven weeks following initial immunization, with an increase in vaccine cargo dose being predictive of more robust protection. Day 1 signatures are inversely correlated with viral loads measured in the airways. *N* = 25 macaques per group. A Spearman-rank test was used to calculate correlations, and *p-*values were adjusted for multiple comparisons using a Bonferroni correction. From left to right, signatures are correlated to: **NT50** titers, anti-RBD antibody responses (**ELISA**), T cell activity (**ELISPOT**), and viral loads measured in the bronchoalveolar lavage (**BAL**), and nasal swabs (**NS**).

These early transcriptomic signatures on day 1 also correlated inversely with viral loads (**Figure 4**) from bronchoalveolar lavage (BAL) (−0.70 ≤ ρ ≤ 0.05; 1.1 x 10^−4^ ≤ *p*_adj_ ≤ 8.2 x 10^−1^) and nasal swabs (NS) (0.14 ≤ ρ ≤ 0.67; 2.3 x 10^−4^ ≤ *p*_adj_ ≤ 5.0 x 10^−1^) following SARS-CoV-2 challenge at week 24, suggesting the relevance of the early innate profiles in shaping adaptive immunity and protective efficacy months later (**Figure 4**). We observed significant and positive correlations of antiviral signature (*ANTIVIRAL IFN SIGNATURE*) with NT50 titers (ρ = 0.72, *p*_adj_ = 5 x 10^−5^) and ELISA antibody titers (ρ = 0.74, *p*_adj_ = 3 x 10^−5^) at day 42 post-prime. Conversely, interferon activity correlated negatively with viral loads in both BAL (ρ = −0.34, *p*_adj_ = 0.1) and NS (ρ = −0.54, *p*_adj_ = 5 x 10^−3^) on day 1 post-challenge (day 43 post-prime). Correlations with ELISpot cellular activity remain significant until day 28 post-prime (ρ = 0.67, *p*_adj_ = 2 x 10^−4^) (**Figure 4**). This suggest that initial innate immune pathways correlate with vaccine protective efficacy^21^.

### Transcriptomic pathways on day 7 shaped adaptive immune responses

We next performed ssGSEA^18^ and Spearman rank correlations of transcriptomic pathways on day 7 with adaptive immune responses (**Supplementary Table 4**). We found that CD4^+^ T cell (0.02 ≤ ρ ≤ 0.52; 8.4 x 10^−3^ ≤ *p*_adj_ ≤ 9.4 x 10^−1^) and mitotic cell cycle pathways (−0.05 ≤ ρ ≤ 0.66; 2.9 x 10^−4^ ≤ *p*_adj_ ≤ 8.1 x 10^−1^) correlated positively with ELISA, NT50, and ELISpot data on day 42 (**Figure 5**). Conversely, they correlated negatively with viral loads in BAL (−0.48 ≤ ρ ≤ 0.22; 1.4 x 10^−2^ ≤ *p*_adj_ ≤ 9.4 x 10^−1^) and NS (−0.46 ≤ ρ ≤ -0.06; 1.9 x 10^−2^ ≤ *p*_adj_ ≤ 7.9 x 10^−1^) following SARS-CoV-2 challenge (**Figure 5**). These data are consistent with a previous study that showed that pre-boost CD4^+^ T cell numbers correlated with post-boost CD8^+^ T cell numbers and antibody levels^22^. While CD4^+^ T cell pathways and mitotic cell cycle pathways on day 7 correlated with adaptive immunity and protection^23^, extracellular matrix (−0.60 ≤ ρ ≤ 0.20; 1.3 x 10^−3^ ≤ *p*_adj_ ≤ 3.4 x 10^−1^) and integrin cell surface interactions (−0.40 ≤ ρ ≤ 0.04; 4.7 x 10^−2^ ≤ *p*_adj_ ≤ 9.9 x 10^−1^) correlated negatively with immunogenicity and protection. Previous studies have demonstrated that integrin interactions^24^ and extracellular matrix proteins^25^ contribute to SARS-CoV-2 infection.

**Figure 5:**
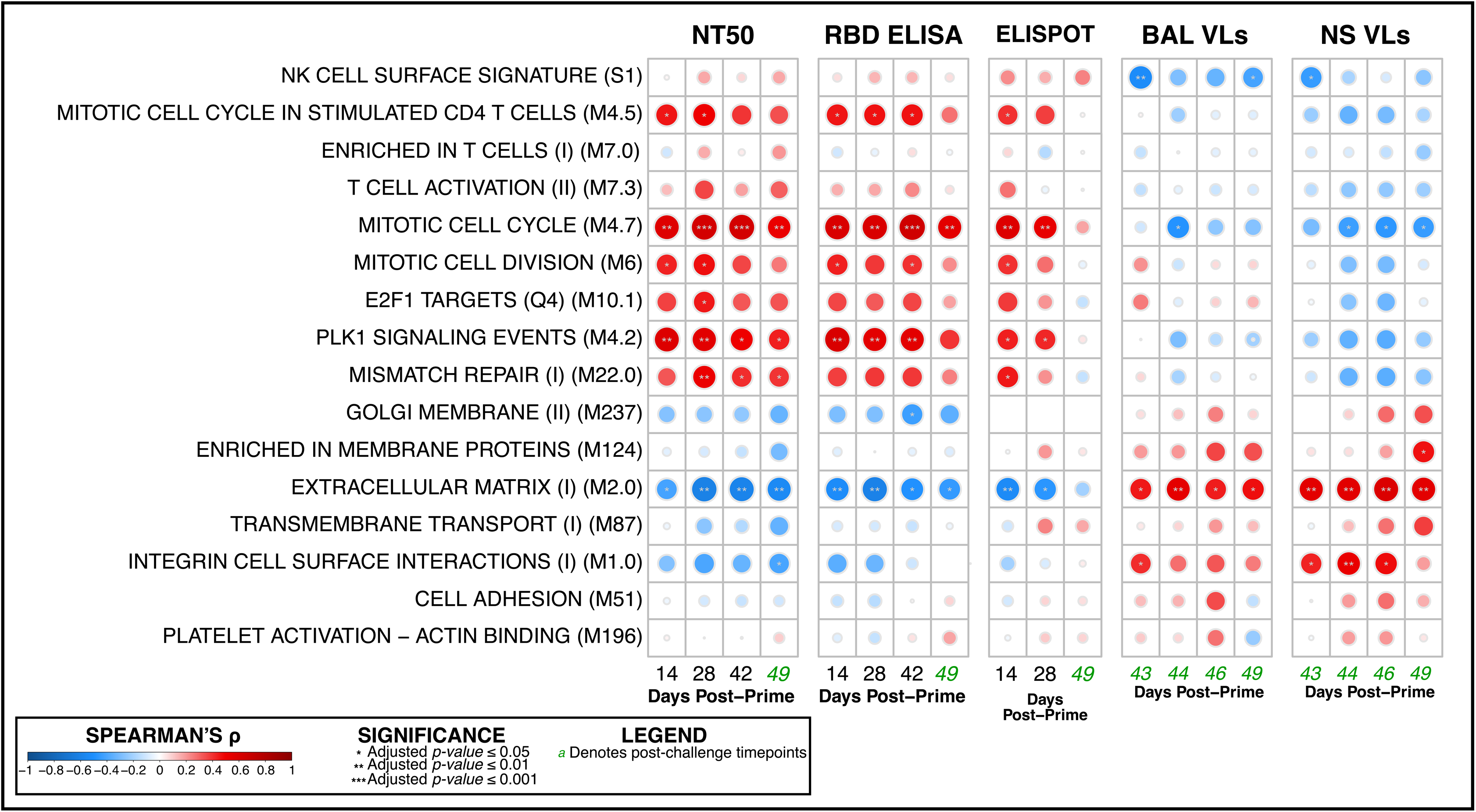
Whole-blood transcriptomic T cell and mitotic signatures at day 7 following initial Ad26.COV2.S immunization in macaques are predictive of antibody and T cell responses at two to seven weeks following initial immunization, with an increase in vaccine cargo dose being predictive of more robust protection. Conversely, pathways associated with extracellular metabolism are inversely correlated with protection, and positively correlated with viral loads in the airways. *N* = 25 macaques per group. A Spearman-rank test was used to calculate correlations, and *p-*values were adjusted for multiple comparisons using a Bonferroni correction. From left to right, signatures are correlated to: **NT50** titers, anti-RBD antibody responses (**ELISA**), T cell activity (**ELISPOT**), and viral loads measured in the bronchoalveolar lavage (**BAL**), and nasal swabs (**NS**).

### Comparable activation of transcriptomic signatures in humans and macaques following Ad26.COV2.S immunization

To characterize the innate pathways induced by Ad26.COV2.S immunization in humans, we performed a whole-blood transcriptomic analysis of samples from clinical trial participants in a phase 1/2a clinical study (NCT04436276)^7^ (**Figure 1B**). The study evaluated the safety and immunogenicity of the in healthy adults and included assessments of humoral and cellular immune responses^7^. Peripheral blood was drawn and bulk RNA sequencing was performed for 20 participants who received 1 x 10^11^ vp (*n* = 10) or 5 x 10^10^ vp (*n* = 10) Ad26.COV2.S at day 1 and day 7 following immunization^8^. Using GSEA^12, 13, 26^, we performed pathway analysis and observed that the key innate immune pathways observed following Ad26.COV2.S immunization of macaques (**Figure 2**) were recapitulated in humans (**Figure 6, Supplementary Table 5**), including interferon activity (*ANTIVIRAL IFN SIGNATURE*, NES = 2.74, FDR = 4.94 x 10^−5^), TLR activity (*TLR AND INFLAMMATORY SIGNALING*, NES = 2.47, FDR = 4.94 x 10^−5^), dendritic cells (*ENRICHED IN ACTIVATED DENDRITIC CELLS*, NES = 2.84 FDR = 4.94 x 10^−5^) and monocytes (*ENRICHED IN MONOCYTES*, NES = 3.10 FDR = 4.94 x 10^−5^), as well as acute inflammatory signals (*CHEMOKINES AND INFLAMMATORY MOLECULES IN MYELOID CELLS*, NES = 1.73 FDR = 2.32 x 10^−2^) on day 1 and T cell cycle activity on day 7 (*MITOTIC CELL CYCLE IN STIMULATED CD4 T CELLS*, NES = 2.42, FDR = 4.94 x 10^−5^). Similarly, an MDS plot for our participant data shows even more distinct clustering for day 1 samples, as compared to macaques (**Figures 1C, 1D**)^11^. These data show that innate immune responses following SARS-CoV-2 infection in humans are similar to macaques^27^, which suggests the comparability of these two models^20^. In addition to corroborating our results in **Figure 2**, we observed that our pathway analysis in **Figure 6** agrees with our results from our previous study of the same cohort of participants^8^. For example, we observed consistently robust interferon activity in participants who received a dose of 1 x 10^11^ vp (*ANTIVIRAL IFN SIGNATURE*, NES = 2.70, FDR = 4.94 x 10^−5^) and those who received a dose of 5 x 10^10^ vp (*ANTIVIRAL IFN SIGNATURE*, NES = 2.53, FDR = 1.14 x 10^−4^).

**Figure 6:**
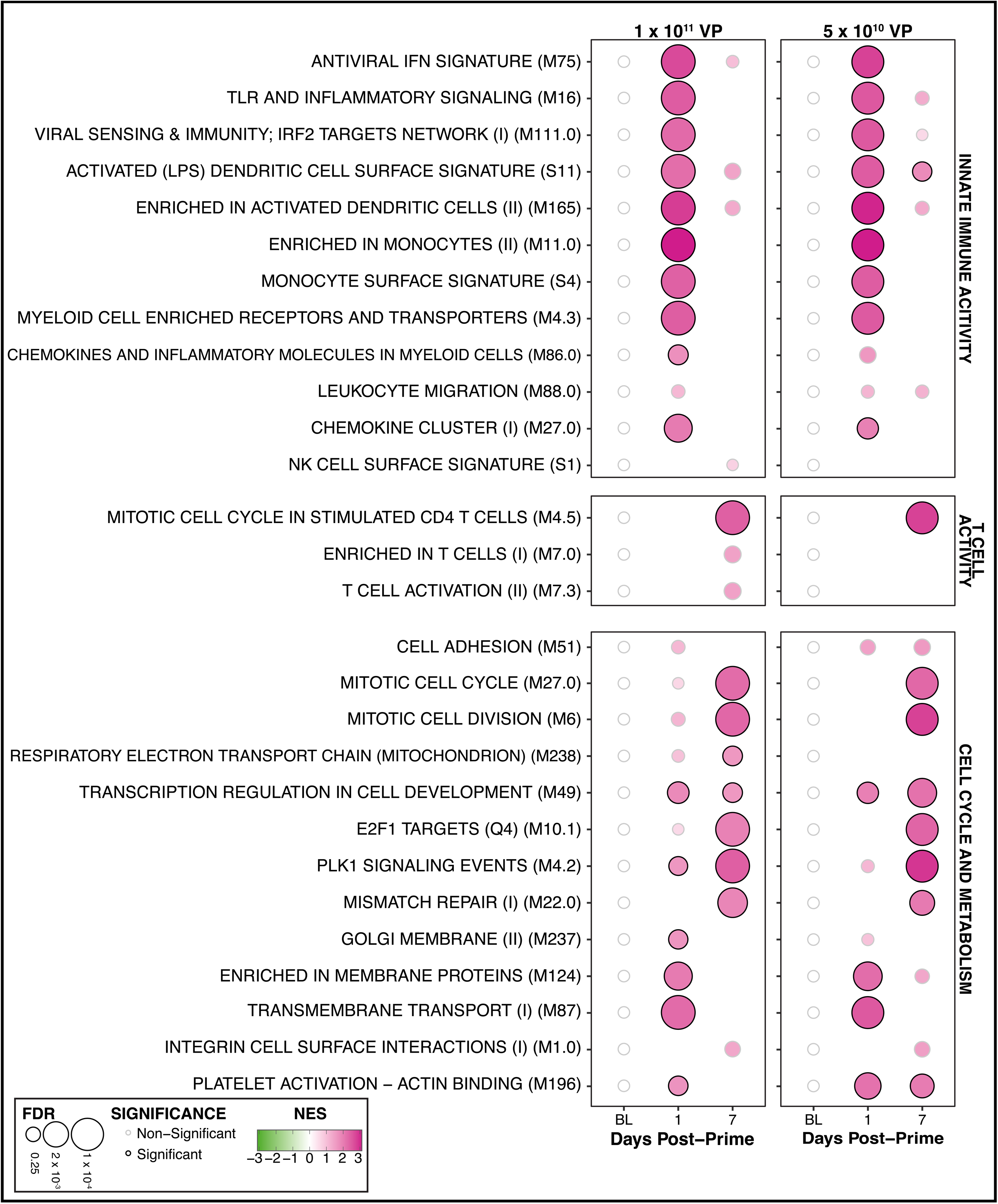
Whole-blood transcriptomic innate immune signatures at day 1 following initial Ad26.COV2.S immunization in clinical trial participants exhibit similar expression patterns to pathway activation demonstrated in macaques.

### Transcriptomic pathway analysis in humans revealed day 1 activation of innate immune pathways that shaped adaptive immune responses

We previously reported that antibody responses following Ad26.COV2.S immunization of humans persisted for over 238 days^6^. To assess whether day 1 innate immune signatures shaped adaptive immune response in humans as they did in macaques, we calculated individual participant pathway enrichment scores using ssGSEA^18^, and correlated them to immunological outcomes at days 28 and to 238 against three different SARS-CoV-2 variants: WA1/20 (original Wuhan strain), Delta (B.1.617.2), and Omicron (B.1.1.529)^28^. Bonferroni-corrected Spearman-rank tests were performed between ssGSEA scores and the following outcomes: CD4^+^ and CD8^+^ T-cell responses measured by peptide-stimulated intracellular staining assays (ICS), anti-RBD ELISA responses, IFN-γ ELISpot T cell activity, as well as plasma neutralizing antibody (NAb) titers (**Figure 7**). Both WA1/20 and Delta variants showed a robust and significant correlation between innate immune pathways and adaptive immune responses. Specifically, antiviral interferon signatures (0.26 ≤ ρ ≤ 0.59; 1.7 x 10^−3^ ≤ *p*_adj_ ≤ 2.2 x 10^−1^), complement activation (0.19 ≤ ρ ≤ 0.58; 2.2 x 10^−3^ ≤ *p*_adj_ ≤ 4.3 x 10^−1^), TLR and viral sensing pathways (0.23 ≤ ρ ≤ 0.64; 5.3 x 10^−4^ ≤ *p*_adj_ ≤ 3.4 x 10^−1^), as well as inflammatory chemokine modules (0.09 ≤ ρ ≤ 0.64; 5.2 x 10^−4^ ≤ *p*_adj_ ≤ 6.9 x 10^−1^) correlated strongly to adaptive immune responses — particularly to CD8+ T cell activity and anti-RBD activity — at day 28 following initial immunization. Correlations against Omicron responses reduced in magnitude and significance at day 28 for all pathways (**Figure 7**).

**Figure 7:**
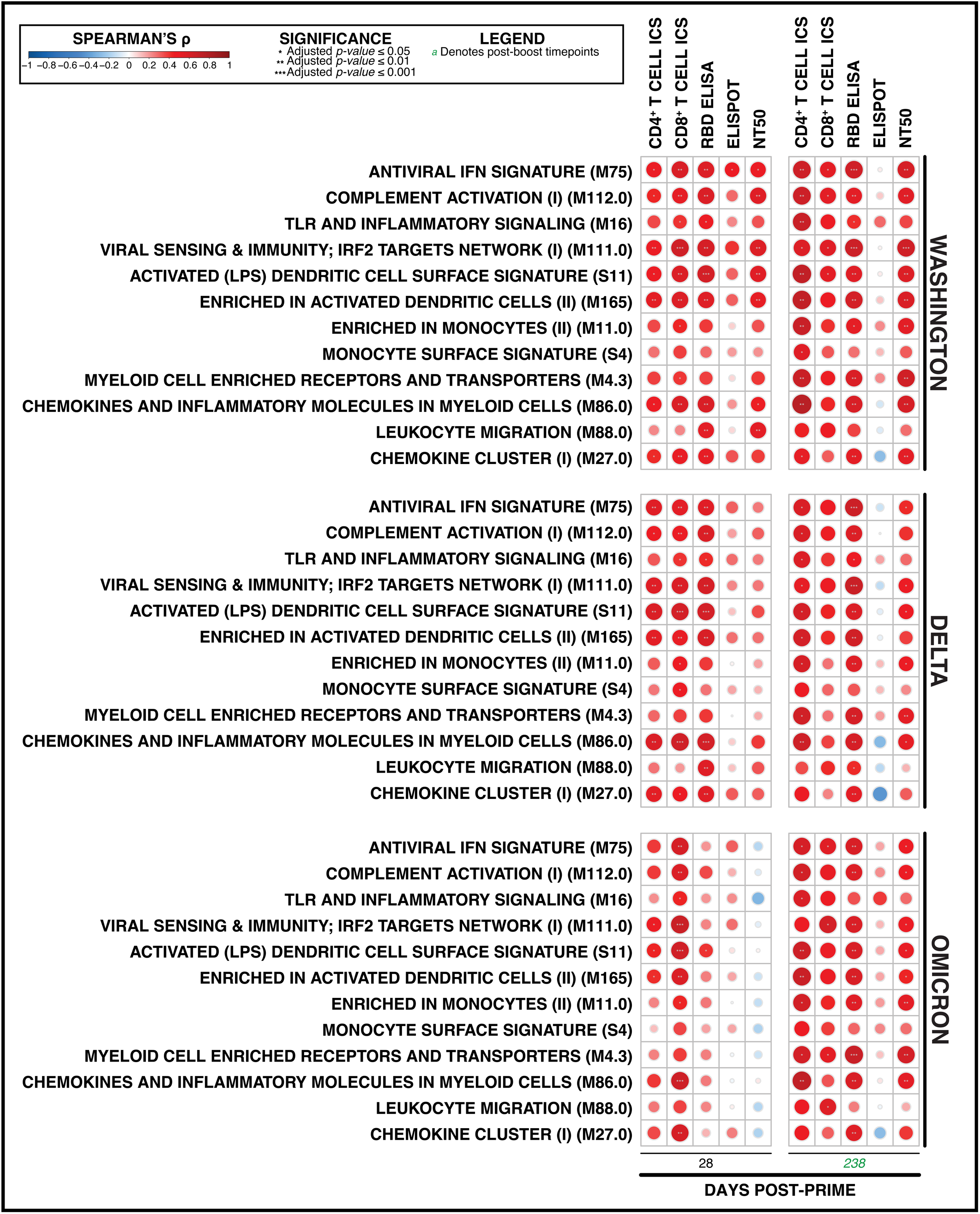
Whole-blood innate immune transcriptomic signatures at day 1 following initial Ad26.COV2.S immunization in participants are predictive of antibody and T cell responses at 28 and 238 days following initial immunization, with an increase in vaccine cargo dose being predictive of more robust protection. A Spearman-rank test was used to calculate correlations, and *p-*values were adjusted for multiple comparisons using a Bonferroni correction. From left to right, signatures are correlated to: CD4+ T cell activity measured by intracellular staining (**ICS**), CD8+ T cell activity measured by **ICS**, anti-RBD antibody responses (**ELISA**), T cell activity (**ELISPOT**), and **NT50** titers.

At day 238 following initial immunization, correlation patterns remained consistent with day 28 activity, with the Omicron strain showing a slight increase in breadth and significance (**Figure 7**). To quantify the effect of the booster dose (administered at day 56 post-prime), we performed Spearman-rank tests to correlate ssGSEA at day 57 post-prime (day 1 post-boost) to adaptive immune responses at day 238 post-prime. For all three variants, innate immune signatures activated by the booster dose seemed to be predictive of CD4+ T cell responses (**Supplementary Figure 1**). This suggested that the innate immune signatures such as antiviral interferon activity on day 1 following initial immunization were the strongest correlates of immunogenicity. Pathways induced at day 7 post-prime (**Figure 8**) and day 7 post-boost (**Supplementary Figure 2**) showed weaker correlations. Statistics are presented in **Supplementary Table 6**. Taken together, our pathway enrichment analysis suggest that the innate immune pathways induced at day 1 following initial Ad26.COV2.S immunization in humans were the strongest predictor of robust adaptive immune responses, which persisted up to approximately 8 months following initial immunization.

**Figure 8:**
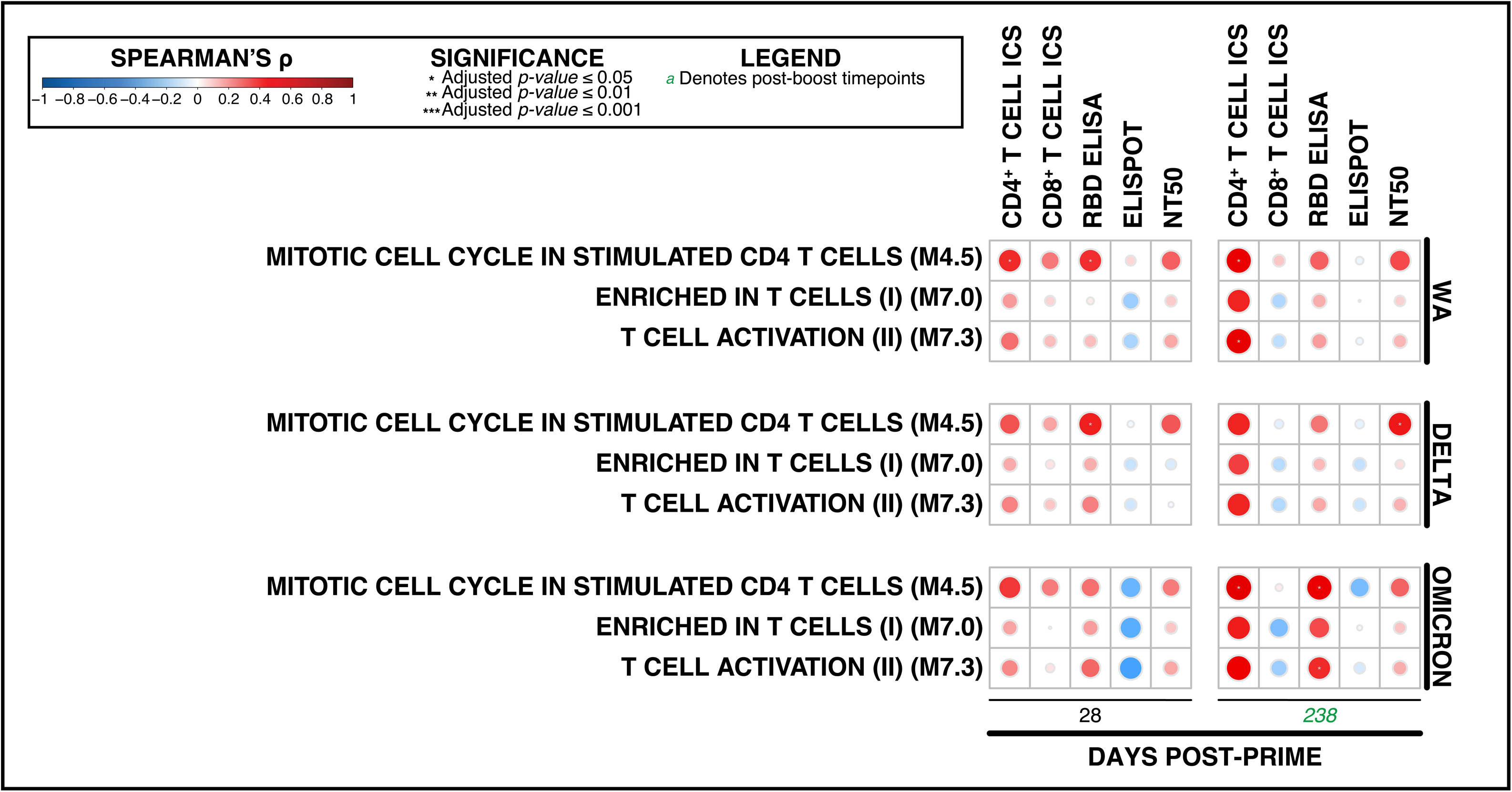
Whole-blood innate immune transcriptomic signatures at day 7 following initial Ad26.COV2.S immunization in participants demonstrate mild yet unsignificant predictive ability of antibody and T cell responses at 28 and 238 days following initial immunization. A Spearman-rank test was used to calculate correlations, and *p-*values were adjusted for multiple comparisons using a Bonferroni correction. From left to right, signatures are correlated to: CD4+ T cell activity measured by intracellular staining (**ICS**), CD8+ T cell activity measured by **ICS**, anti-RBD antibody responses (**ELISA**), T cell activity (**ELISPOT**), and **NT50** titers.

### Consistent activation of proteomic signatures in humans and macaques following Ad26.COV2.S immunization

To complement the transcriptomic signatures, we performed proteomic profiling^29^ using day 1 post-prime plasma from NHP (*n* = 5) and humans (*n* = 10), both immunized with 5 x 10^10^ VP Ad26.COV2.S, using the SomaScan platform. Pathway analysis was performed using the over-representation analysis in g:Profiler’s gGOSt using the Reactome reference database^30, 31^ (**Supplementary Table 7**). We observed significant activity (FDR ≤ 0.5) for antiviral and innate immune pathways and cytokine and interleukin signaling pathways activated in both humans (5.64 x 10^−8^ ≤ FDR ≤ 4.2 x 10^−2^) and macaques (7.24 x 10^−3^ ≤ FDR ≤ 4.9 x 10^−2^) (**Figure 9**). Additionally, intersectional differential gene expression reveals four proinflammatory and innate immune genes upregulated in both transcriptomics and proteomics, for human and macaque samples: *CXCL10, CXCL11, CD274,* and *TLR5* (**Figure 10**).

**Figure 9:**
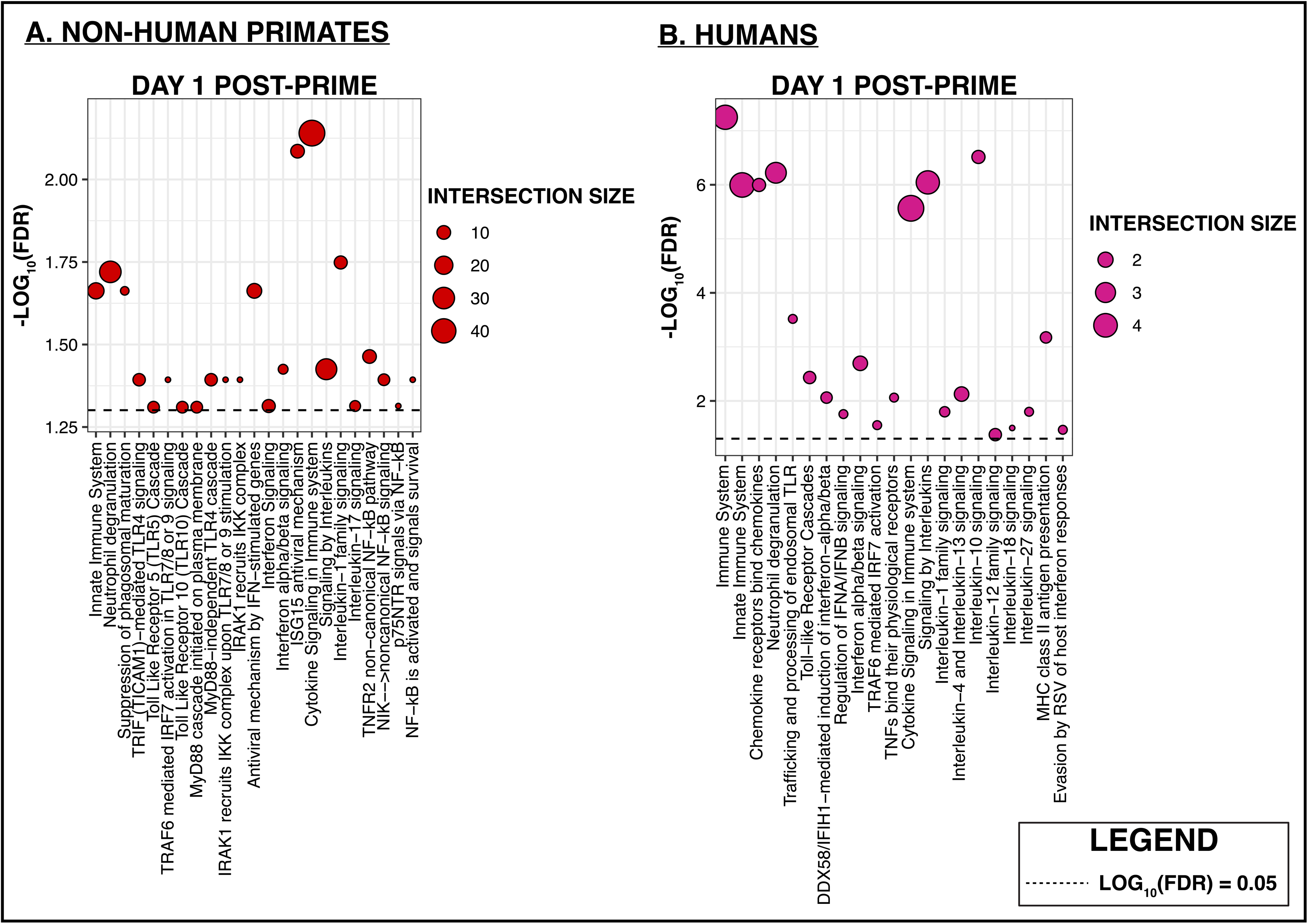
Plasma proteomic signatures reveal robust day 1 post-vaccination pathways. **A.** gProfiler results spanning Reactome innate immune pathways are shown for NHP samples. **B.** gProfiler results spanning Reactome innate immune pathways are shown for participant samples.

**Figure 10:**
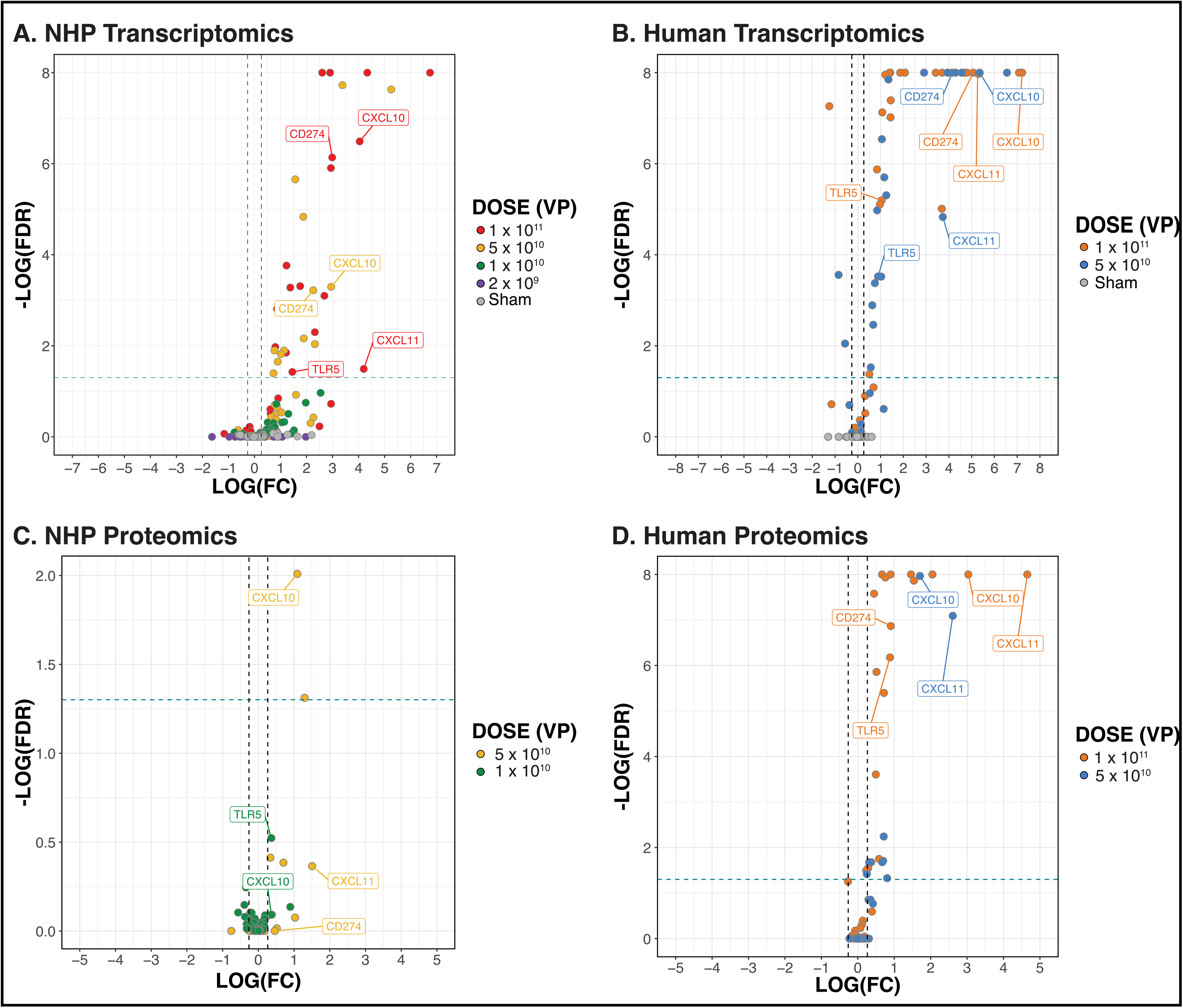
Volcano plot showing differentially expressed genes plotted as a function of their log-fold change from baseline to day 1 post-prime, as well as their FDR.

## DISCUSSION

Ad vectors have been utilized extensively for vaccines and gene therapy, including for multiple COVID-19 vaccines during the pandemic. Moreover, an Ad-HPV vector has recently been approved by the FDA as immunotherapy for recurrent respiratory papillomatosis (Papzimeos; zopapogene imadenovec-drba)^32^. However, innate correlates of immunogenicity and protection efficacy have not yet been fully described^9^. In this study, we analyzed transcriptomic and proteomic signatures in non-human primates and humans following vaccination with Ad26.COV2.S and defined early innate immune pathways that correlated with adaptive immune responses^9^.

We observed that innate immune pathways such as including antiviral, interferon, cytokine/chemokine, dendritic cell, and acute inflammatory responses peaked on day 1 following vaccination, consistent with early priming of the adaptive immune response, and subsequently returned to baseline. By day 7, we detected a marked increase in signatures associated with T and B cell signaling, cell cycle, and metabolic pathways, indicating robust activation of the adaptive immune response. Both phases correlate with downstream immunogenicity and protection in rhesus macaques.

Our data aligns with previous findings in humans, mice and macaques^10, 14^. In mice, we previously demonstrated that an immunization with Ad26 induced innate immune pathways as early as one hour following immunization, with resolution by day 3^14^. In the present study, we observed a similar rapid induction of innate immune pathways at day 1 in both humans, and macaques, with resolution largely occurring by day 3 (**Figure 2**).

In addition, the present results align with our findings that showed robust, dose-dependent activation on day 1 innate immune pathways in participants who received both 1 x 10^11^ vp, and 5 x 10^10^ vp Ad26.COV2.S^8^. Our findings suggest a model in which there is a stereotypical and sequential program that is initiated immediately after immunization by activation of interferon signaling, inflammatory pathways, and immune cell recruitment and leads to a cascade that results in induction of protective adaptive immune responses. Moreover, our data suggest the potential that very early biomarkers may be predictive of long-term vaccine protection.

Given the strong association between DC activity and PLK1 signaling (**Figure 4)**, we can posit that Ad26.COV2.S spike may trigger a TLR-based, DC-mediated innate immune response, with high antiviral interferon activity, necessary components for the activation and proliferation of CD4+ and CD+ T cells as well as anti-SARS-CoV-2 antibodies^33^. We also observed *CXCL10* and *CXCL11* (**Figure 10**), which have been associated with increased immune activity in COVID-19 patients and were previously reported to help shape adaptive and the proliferation and differentiation of T lymphocytes^34, 35^. While TLR5 is classically associated in the recognition of bacterial flagella, emerging evidence has shown that it heterodimerizes with TLR4, and biases the latter towards Myd88-dependent activation of the immune system^36^. TLR4 has been previously shown to be activated by SARS-CoV-2 spike protein^37^. CD274 has been shown to be upregulated in COVID-19 but may serve as an indicator of disease severity^38^. Our findings align well with previous systems vaccinology studies investigating SARS-CoV-2 and other pathogens. Hagan et al. integrated multi-omic and immunological profiling from over 800 patients vaccinated with 13 different vaccines, including HIV and yellow fever, and observed a largely conserved activation pattern of innate immune pathways at day 1 following immunization^39^. Similarly, Nakaya et al. analyzed transcriptomic data from five different seasonal influenza vaccines and identified conserved interferon and DC signatures (across multiple influenza seasons) predictive of durable antibody responses^40^. These and other studies provide a foundational framework that our study builds upon. Effective vaccine design is predicated on eliciting durable protection, and systems vaccinology remains a fundamental tool in elucidating molecular and cellular factors that contribute to vaccine efficacy and protectivity immunity^1, 41^.

In conclusion, we demonstrated that Ad26.COV2.S immunization in rhesus macaques and humans results in rapid induction of antiviral and acute inflammatory and innate immune responses by day 1 that led to subsequent activation of cellular immune pathways by day 7 and that correlates with the magnitude of antigen-specific humoral and cellular immune responses. These data suggest that early innate responses shape adaptive immune responses in a stereotypic fashion, with implications for understanding mechanisms of vaccine protection.

### Limitations of Study

Future investigations should include transcriptomic profiling of animals or humans who received an Ad26 vector with no insert to confirm which pathways are vector-mediated or transgene-mediated^42^. Additional insights could be gained by employing single-cell sequencing in tissues targeted during infection^43^. Additionally, future studies could profile participants following SARS-CoV-2 infection^6–8^. Lastly, pre-existing immunity to homologous coronaviruses has been shown to affect disease phenotype and vaccine response in COVID-19^44^. Thus, pre-existing immunity to coronaviruses should be taken into account, including for the design of a pan-sarbecovirus vaccine^44, 45^.

## AUTHOR CONTRIBUTIONS

D.H.B., K.E.S., F.W., R.C.Z., M.A., and A.C. designed the study. Transcriptomic and proteomic analyses were performed by A.C. Clinical trials were conducted by D.H.B., K.E.S. and A.Y.C. The Ad26 were provided by F.W. and R.C.Z. The paper was written with D.H.B., M.A and A.C. and all co-authors.

## ACKNOWLEDGEMENTS

We thank C. Ayad, G. Dagotto, S. Mohammadabi, F. Mostefai, and A. Rössler, for generous advice, and assistance. We acknowledge support from the NIH (CA260476) and the Massachusetts Consortium for Pathogen Readiness. We thank the participants and staff at the Center for Virology and Vaccine Research Clinical Trials Unit and the Harvard Catalyst Clinical Research Center.

## DATA AVAILABILITY

Raw data for the bulk RNA sequencing studies are available in NCBI GEO under the accession number GSE220659. All other supporting data are available upon request from the corresponding author.

## MATERIALS & METHODS

### Cohorts

#### Animal Cohort

The study design has been previously been described ^10^. Thirty outbred Indian-origin adult male (10) and female (20) rhesus macaques (*Macaca mulatta*) were randomly allocated to groups. Animals received a single immunization of 1×10^11^, 5×10^10^, 1.125×10^10^, or 2×10^9^ viral particles (vp) Ad26.COV2.S (Janssen; n = 5/group) or sham (n = 10) by the intramuscular route without adjuvant at week 0. At week 6, all animals were challenged with 1.0×10^5^ TCID50 (1.2×10^8^ RNA copies, 1.1×10^4^ PFU) SARS-CoV-2, which was derived from USA-WA1/2020 (NR-52281; BEI Resources) and deep sequenced. Animal studies were conducted in compliance with all relevant local, state, and federal regulations and were approved by the Bioqual Institutional Animal Care and Use Committee (IACUC) ^10^.

#### Human Cohort

Samples were collected from participants of a clinical trial, which has been previously described ^7^. This study was conducted at a single site at Beth Israel Deaconess Medical Center in Boston, and was approved by the hospital’s institutional review board. Briefly, this study was performed to evaluate the safety, reactogenicity, and immunogenicity of Ad26.COV2.S at 5 × 10^10^ or 1 × 10^11^ viral particles administered intramuscularly as single-shot or 2-shot vaccine schedules, 56 days apart, in healthy adults. The cohort described here enrolled adults 18 to 55 years of age (N = 25), and included visits and sampling of serum, plasma, and peripheral blood mononuclear cells to allow for investigation of exploratory end points ^7^.

### Bulk RNA Sequencing

#### Animal Cohorts

Buk RNA sequencing was collected, quality controlled, and read counts were generated as previously described ^8^. Total RNA was extracted from whole blood samples using the PAXgene Blood RNA Kit IVD (PreAnalytix). RNA quality was assessed with the Fragment Analyzer (Agilent) and its Standard Sensitivity RNA kit. Total RNA was normalized to 100 ng before random hexamer priming and libraries were generated by the TruSeq Stranded Total RNA—Globin kit (Illumina). The resulting libraries were assessed on the Fragment Analyzer (Agilent) with the High Sense Large Fragment kit and quantified using a Qubit 3.0 fluorometer (Life Technologies). Medium-depth sequencing, with at least 30 million reads per sample, was performed on a NovaSeq 6000 (Illumina) with a PE100 configuration, paired-end run yielding ∼40 million reads per sample.

#### Human Cohorts

Peripheral blood mononuclear cell (PBMC) were collected from the Ad26.COV2.S vaccinated individuals as PAXgene Blood samples were not available for the Ad26.COV2.S vaccinated individuals. Cryopreserved PBMC samples were thawed, counted, and assessed for viability, then 1 million PBMCs were pelleted and lysed in 350 uL of RLT with beta-mercaptoethanol. RNA was isolated using the RNeasy Mini kit (Qiagen). RNA quality was assessed with the Fragment Analyzer (Agilent) and its Standard Sensitivity RNA kit. Total RNA was used to prepare libraries using the TruSeq Stranded Total RNA Globin kit (Illumina). The resulting libraries were assessed on the Fragment Analyzer (Agilent) with the High Sense Large DNA Fragment kit and quantified using a Qubit 3.0 fluorometer (Life Technologies). Medium to high-depth sequencing (> 40 million reads per sample) was performed on the NovaSeq 6000 platform (Illumina) with a 100-base pair, paired-end run design ^8^.

#### Quality Control and Abundance Estimates

The quality of the raw reads was assessed using multiQC^46^. Raw demultiplexed fastq paired-end read files were trimmed of adapters and filtered using the program skewer to discard those with an average phred quality score of less than 30 or a length of less than 36. Human-sequenced data were aligned to the Homo Sapiens NCBI reference genome assembly version GRCh38, and the rhesus macaques data were aligned to the Mmul10-100 and the MacaM assemblies and annotations of the Indian rhesus macaque genome using STAR version 2.7.3a ^47^. Transcript abundance estimates were calculated internal to the STAR aligner using the algorithm of htseq-count.

### SomaLogic Proteomics

Collection of SomaLogic proteomics data has been previously described by Aid et al.^8^ Briefly, 55 μl of plasma from participants and animals, five pooled serum controls, and one buffer control were analyzed using the SomaScan Assay Kit for human serum V4.1 (Cat#. 900-00021), measuring the expression of 6596 unique human protein targets using 7596 SOMAmer (slow off-rate modified aptamer) reagents, single-stranded DNA aptamers, according to the manufacturer’s standard protocol (SomaLogic; Boulder, CO). The assay used standard controls, including 12 hybridization normalization control sequences used to control for variability in the Agilent microarray readout process, as well as five human calibrator control pooled serum replicates and 3 Quality Control (QC) pooled replicates used to mitigate batch effects and verify the quality of the assay run using standard acceptance criteria. The readout is performed using Agilent microarray hybridization, scan, and feature extraction technology. Normalization and calibration were performed using internal SomaLogic controls and references^8, 48^.

### Bioinformatics Analyses

All analyses were performed locally using R 4.4.2^49^. Functions were used with default parameters unless noted otherwise.

### Bulk RNA Sequencing

#### Batch Correction

Batch correction with the sva package’s ComBat_seq() function was performed on rhesus macaque bulk RNA-sequencing counts, as the total abundance matrix comprised of two different animal studies ^50^.

#### Raw Read Count Normalization

Raw read counts were normalized using the edgeR package (version 3.2.0) ^51^. Briefly, batch corrected raw counts were transformed into log-normalized read counts for further downstream analysis. Library scaling was performed with the calcNormFactors() function, with the method set to the Trimmed Mean of M-values (method = “TMM”) option. Log counts-per-million were then estimated using the cpm() function with the log flag set to true (log = TRUE) ^51^.

#### Pathway Analysis

Pathway analysis was performed with Gene Set Enrichment Analysis (GSEA) ^12^, version 4.2.3. Our input expression data set were the normalized counts generated with edgeR^51^. Genes in the expression data had been previously annotated and non-human primate genes were previously mapped to their human orthologs^8, 52^. As such, we set the Collapse/Remap option to “No_Collapse”. We ran GSEA^12^ against a set of pathways suitable for whole-blood transcriptomic analysis: the Blood Transcription Modules (BTM) dataset generated by Li et al.^13^ For each dose, in both animal, and human cohorts, each post-vaccination timepoint was contrasted to baseline/pre-vaccination values, and the appropriate phenotype label file was supplied for each run. Because our discovery (NHP) cohort had fewer than seven samples per group, we used the “genotype” permutation type to run GSEA. To retain consistency, we also used the “genotype” permutation type in our validation (human) cohort^12^. As such, only the pathways that had a False Discovery Rate (FDR) ≤ 0.05 were deemed significant, as per the GSEA manual^12^. Then, only significant pathways were considered for further analysis.

In the non-human primate cohort, day 1 post-prime pathways in the sham group that were significantly modulated compared to baseline were excluded from analysis in order to specifically gauge vaccine-induced pathways. Moreover, to account for inherent variability at baseline between vaccinated and sham animals, we contrasted day 0 expression data in the high-dose (1 x 10^11^ vp) group to day 0 expression data in the sham group, and excluded any significantly induced pathways from downstream analysis. Longitudinal pathway activity was visualized with dotplots generated with ggplot2, version 3.4.4^53^.

#### Correlation Analysis

For the rhesus macaque datasets, normalized enrichment scores (NES) were estimated for each animal using single sample GSEA (ssGSEA) from the GSVA package^18^ in R^49^. The NES returned by ssGSEA were then correlated with immunological outcome measurements described above, namely: neutralizing antibody titers, S and RBD antibody titers, and IFN-γ mediated ELISPOT SFCs. This was paired with correlations to post-challenge subgenomic mRNA of SARS-COV-2 in both nasal swabs and bronchoalveolar lavage. Spearman rank tests were performed and we adjusted p-values using the Bonferroni correction for multiple testing in the corr.test() function from the psych package^54^. Correlation matrices were visualized with the corrplot package^55^.

#### Differential Gene Expression

Raw read counts were normalized to log counts-per-million using edgeR’s Trimmed Mean of M-values. Differential gene expression (DGE) was then performed with the limma package^56^, version 3.64.3, in R^49^. We established a design matrix using the model.matrix function with “∼Time” being our formula. A linear model for each gene is then fit with lmFit function^56, 57^. To assess differential gene expression between day 1 (D1) following immunization and baseline (D0), we used the makeContrasts function in limma^56^, with the contrasts argument set to “TimeD1 – TimeD0”, and the levels argument set to “colnames(coef(fit))’ where fit is the fit returned by the lmFit function^49, 56^. We then used the contrasts.fit to estimate the difference in expression for each gene^56, 57^, and the eBayes functions to shrink errors in our fitted models^56, 57^. We then used the topTable function in limma to generate our list of differentially expressed genes (DEG)^56, 57^. DEGs were then ranked based on the product of their computed log-fold change and the negative log10 of their p-value.

### Plasma Proteomics

#### Differential Gene Expression

Normalized relative fluorescence units (RFUs) provided by SomagLogic^8, 48^ were first log-transformed. Differential gene expression (DGE) was then performed with the limma package^56^, version 3.64.3, in R^49^. We established a design matrix using the model.matrix function with “∼Time” being our formula. A linear model for each gene is then fit with lmFit function^56, 57^. To assess differential gene expression between day 1 (D1) following immunization and baseline (D0), we used the makeContrasts function in limma^56^, with the contrasts argument set to “TimeD1 – TimeD0”, and the levels argument set to “colnames(coef(fit))’ where fit is the fit returned by the lmFit function^49, 56^. We then used the contrasts.fit to estimate the difference in expression for each gene^56, 57^, and the eBayes functions to shrink errors in our fitted models^56, 57^. We then used the topTable function in limma to generate our list of differentially expressed genes (DEG)^56, 57^. DEGs were then ranked based on the product of their computed log-fold change and the negative log10 of their p-value.

#### Pathway Analysis

The ordered list of gene identifiers were submitted to g:Profiler’s gGOSt profiling tool for pathway analysis^30^. Options were tailored to suit or analysis. “Organism” was left to *Homo sapiens* (macaque gene identifiers were preemptively converted to their human orthologs), and we selected only the “Ordered query” option from the initial option menu. In the “Advanced options” menu, we changed the Significance threshold to ‘Benjamini-Hochberg FDR” and left every other parameter to its default setting. In the “Data sources” menu, we selected only the “Reactome” option from the “biological pathways” header^30^. Our readouts were exported as CSVs^30^, imported into R^49^, and visualized with ggplot2^53^.

---

**Figure S1:**
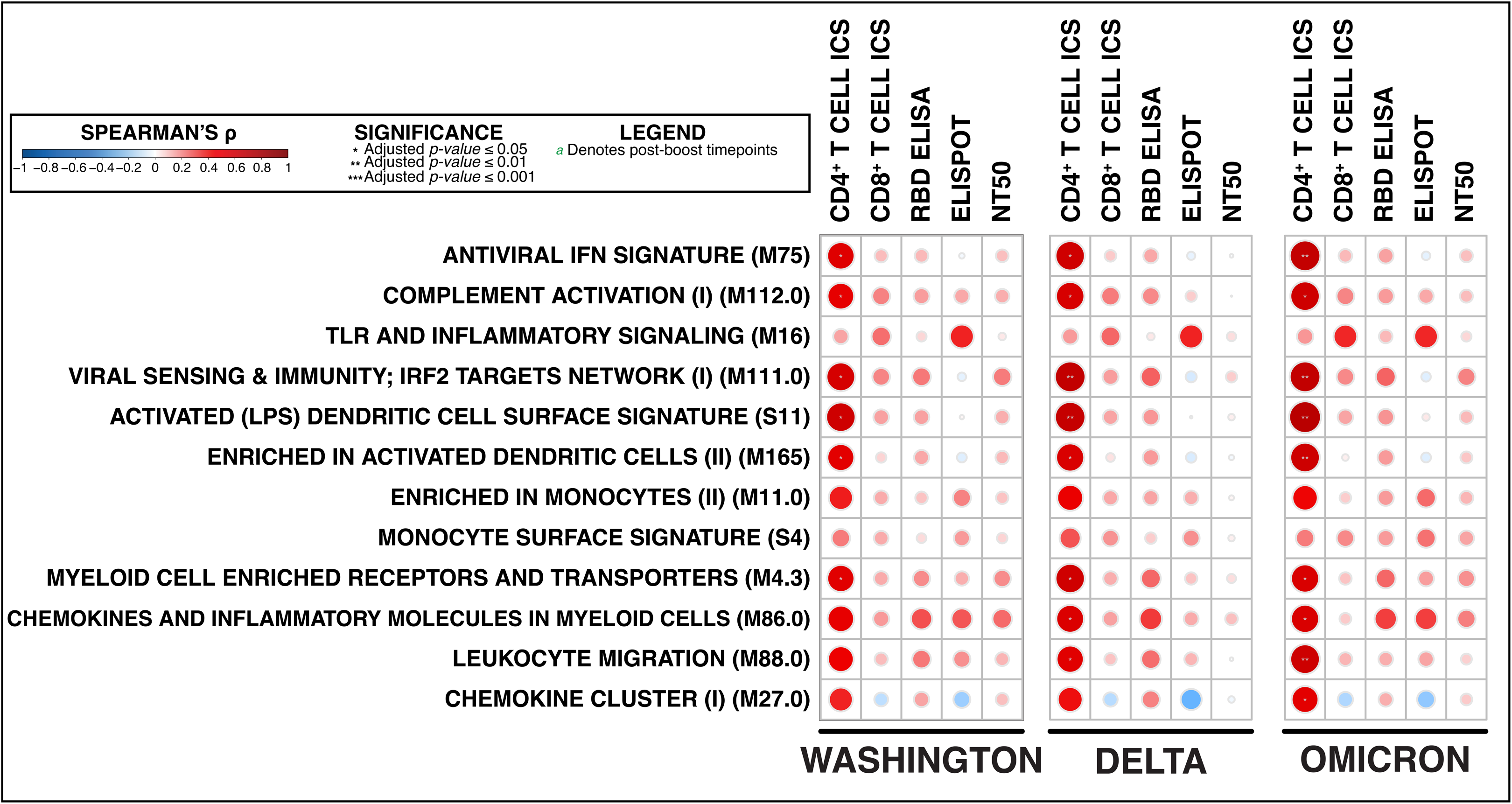
Whole-blood innate immune transcriptomic signatures at day 1 following Ad26.COV2.S boost in participants demonstrate moderate yet largely unsignificant predictive ability of antibody and T cell responses 238 days following initial immunization. A Spearman-rank test was used to calculate correlations, and *p-*values were adjusted for multiple comparisons using a Bonferroni correction. From left to right, signatures are correlated to: CD4+ T cell activity measured by intracellular staining (**ICS**), CD8+ T cell activity measured by **ICS**, anti-RBD antibody responses (**ELISA**), T cell activity (**ELISPOT**), and **NT50** titers.

**Figure S2:**
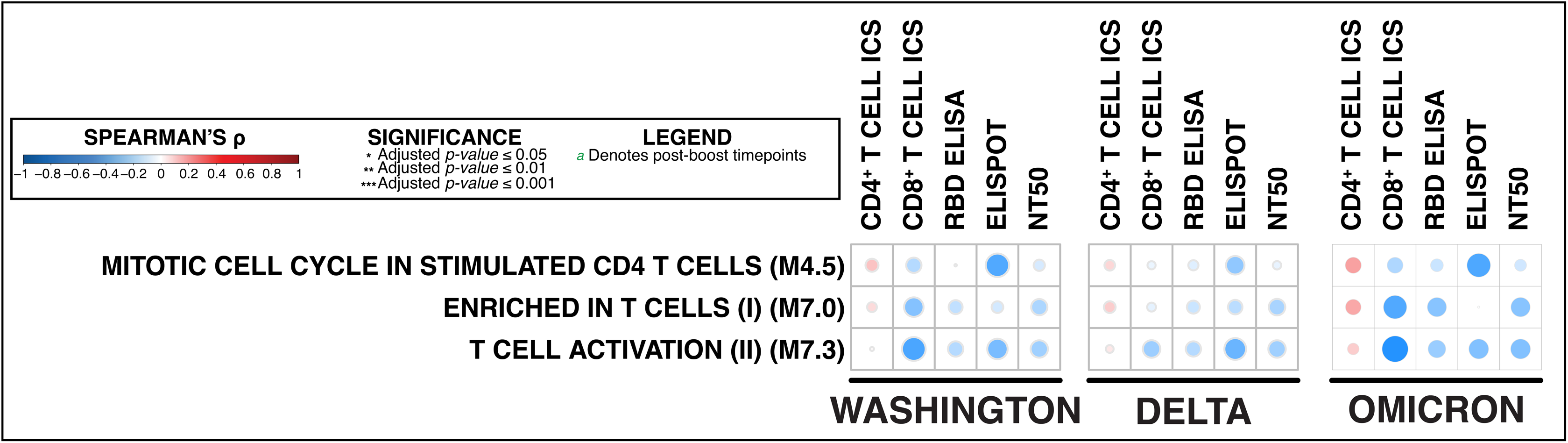
Whole-blood innate immune transcriptomic signatures at day 7 following Ad26.COV2.S boost in participants demonstrate no predictive ability of antibody and T cell responses 238 days following initial immunization. A Spearman-rank test was used to calculate correlations, and *p-*values were adjusted for multiple comparisons using a Bonferroni correction. From left to right, signatures are correlated to: CD4+ T cell activity measured by intracellular staining (**ICS**), CD8+ T cell activity measured by **ICS**, anti-RBD antibody responses (**ELISA**), T cell activity (**ELISPOT**), and **NT50** titers.

